# Amyloidosis and ageing have differential effects on microglial Ca^2+^ activity in the mouse brain

**DOI:** 10.1101/2023.08.10.552670

**Authors:** Pablo Izquierdo, Renaud B. Jolivet, David Attwell, Christian Madry

## Abstract

In microglia, changes in intracellular calcium concentration ([Ca^2+^]_i_) may regulate process motility, inflammasome activation, and phagocytosis. However, while neurons and astrocytes exhibit frequent spontaneous Ca^2+^ activity, microglial Ca^2+^ signals are much rarer and poorly understood. Here, we studied [Ca^2+^]_i_ changes of microglia in acute brain slices using Fluo-4–loaded cells and mice expressing GCaMP5g in microglia. Spontaneous Ca^2+^ transients occurred ∼5 times more frequently in individual microglial processes than in their somata. We assessed whether microglial Ca^2+^ responses change in Alzheimer’s disease (AD) using *App*^NL-G-F^ knock-in mice. Proximity to Aβ plaques strongly affected microglial Ca^2+^ activity. Although spontaneous Ca^2+^ transients were unaffected in microglial processes, they were 5-fold more frequent in microglial somata near Aβ plaques than in wild-type microglia. Microglia away from Aβ plaques in AD mice showed intermediate properties for morphology and Ca^2+^ responses, partly resembling those of wild-type microglia. By contrast, somatic Ca^2+^ responses evoked by tissue damage were less intense in microglia near Aβ plaques than in wild-type microglia, suggesting different mechanisms underlying spontaneous vs. damage-evoked Ca^2+^ signals. Finally, as similar processes occur in neurodegeneration and old age, we studied whether ageing affected microglial [Ca^2+^]_i_. Somatic damage-evoked Ca^2+^ responses were greatly reduced in microglia from old mice, as in the AD mice. In contrast to AD, however, old age did not alter the occurrence of spontaneous Ca^2+^ signals in microglial somata but reduced the rate of events in processes. Thus, we demonstrate distinct compartmentalised Ca^2+^ activity in microglia from healthy, aged and AD-like brains.

## INTRODUCTION

Microglia, the brain-resident parenchymal macrophages, account for up to 12% of brain cells, with a particularly high density in the hippocampus [1]. They control multiple processes including neurogenesis, synapse monitoring and pruning, myelination, vasculogenesis, blood-brain barrier integrity, inflammation, and phagocytosis [2].

Microglial surveillance of the brain parenchyma, chemotactic movement of microglial processes and phagocytosis all depend on actin polymerization and subsequent cytoskeletal rearrangements. These, as well as activation, proliferation and production of inflammatory cytokines, are controlled, at least in part, by Ca^2+^ acting as a second messenger, and rises in [Ca^2+^]_i_ have been reported to be associated with microglial functions including damage-induced chemotactic and phagocytic events [3–7]. In general, phagocytic cells exhibit periodic spontaneous Ca^2+^ transients as well as stimulus-evoked [Ca^2+^]_i_ rises [8], and microglial Ca^2+^ activity is important for the regulation of lysosomes, which process incorporated material [9]. While ‘resting’ microglia rarely exhibit spontaneous Ca^2+^ transients *in vivo*, external pathological stimuli such as damage to nearby neurons or mechanical distortions rapidly raise their [Ca^2+^]_i_ [7, 10].

Previous studies using mainly cultured microglia identified a number of ligands evoking Ca^2+^ responses in microglia, including ATP, ADP and UDP as endogenous stimuli [11–13] and pathology-related substances such as lipopolysaccharide (LPS) and amyloid beta (Aβ) [8]. However, Ca^2+^ responses are significantly different for cultured microglia compared to their counterparts embedded in the native CNS environment [8], as microglia partly lose their characteristic genetic profile under more artificial cell culture conditions [14]. In transgenic mice overexpressing Aβ, accumulation of which is a hallmark of Alzheimer’s disease (AD), altered Ca^2+^ signalling was detected in astrocytes and microglia electroporated with Ca^2+^-sensing dyes [15].

While previous research has relied mainly on Ca^2+^-sensitive dyes, we also took advantage of Cx3cr1^CreER^×GCaMP5g-IRES-tdTomato mice expressing an inducible genetically-encoded Ca^2+^ indicator (GECI) cross-bred with *App*^NL-G-F^ knock-in AD mice [16] to examine microglial Ca^2+^ activity, both spontaneous and upon inducing brain damage, in the presence and absence of AD-related amyloidosis. This strategy allows all cells in the field of view to be captured spatially in relation to plaque pathology and abrogates the need for external manipulation to insert the Ca^2+^ sensor. As similar mechanisms are thought to occur during brain ageing and brain disease [17–22], we also assessed whether the Ca^2+^ activity of microglia was affected differently by AD and normal ageing.

## MATERIALS AND METHODS

### Animal procedures

Procedures were performed in accordance with the UK Animals (Scientific Procedures) Act 1986 (Home Office License 70/8976). Rats (postnatal day 12–14 for work with patch-clamp loading of Ca^2+^-sensitive dye) and mice (postnatal day 120–130 and 300–310 for work with the GECI investigating effects of AD and ageing, respectively) were housed in open-shelf units and individually ventilated cages, respectively, with food and water *ad libitum*. Animals of both sexes were sacrificed by cervical dislocation followed by decapitation (for live imaging), or by an intraperitoneal overdose of pentobarbital sodium (Euthatal, 200 µg/g body weight) followed by transcardial perfusion-fixation with 4% paraformaldehyde (for immunohistochemistry).

For experiments in rats, loading of the Ca^2+^ indicator Fluo-4 was achieved by whole-cell patch-clamping of hippocampal microglia (see below). To study microglial properties in the initial stages of AD-related pathology, 4-month-old homozygous *App*^NL-G-F^ knock-in AD mice or wild-type (WT) littermates were used, where Aβ plaque deposition starts in the AD mice from 2 months of age [16]. For Ca^2+^ imaging, *App*^NL-G-F^ mice were crossed with transgenic mice expressing GCaMP5g-IRES-tdTomato [23] and Cx3cr1^CreER^ [24], to generate offspring where microglia exhibit expression of GCaMP5g and tdTomato following tamoxifen gavage (Sigma T5648; 120 µg/g body weight for four consecutive days). Mice that were homozygous for GCaMP5g, heterozygous for Cx3cr1^CreER^ and either wild-type or homozygous for *App*^NL-G-F^ were used for imaging ≥21 days after the first tamoxifen dose (given at P85–P105).

### Solutions

Acute brain slices (250 µm, parasagittal) containing dorsal hippocampi were prepared on a Leica VT1200S vibratome with ice-cold slicing solution containing (in mM): 124 NaCl, 2.5 KCl, 26 NaHCO_3_, 1 NaH_2_PO_4_, 10 glucose, 1 CaCl_2_, 2 MgCl_2_ and 1 kynurenic acid. Osmolarity was adjusted to ∼295 mOsm/kg and pH set to 7.4 when bubbled with 5% CO_2_/95% O_2_.

For live imaging, brain slices were incubated in artificial cerebrospinal fluid (aCSF) containing (in mM): 124 NaCl, 2.5 KCl, 26 NaHCO3, 1 NaH_2_PO_4_, 10 glucose, 2 CaCl_2_, 1 MgCl, and 0.1 Na-ascorbate. Osmolarity was adjusted to ∼295 mOsm/kg and pH was set to 7.4 with NaOH. Solutions were oxygenated with 20% O_2_/5% CO_2_/75% N_2_.

### Immunohistochemistry

Perfusion-fixed brain slides (250 µm, parasagittal) were prepared on a Leica VT1200S vibratome, permeabilised and blocked for 2 h at room temperature in blocking buffer (10% normal horse serum and 0.02% Triton X-100 in phosphate-buffered saline, PBS), followed by incubation with primary antibodies (mouse anti-human Aβ 82E1 [1:500, IBL 27725], rat anti-CD68 [1:250, BioRad MCA1957], and rabbit anti-Iba1 [1:500, Synaptic Systems 234003]) in blocking buffer for 12 h at 4°C with agitation. Following PBS washes, Alexa-conjugated secondary antibodies (Invitrogen) diluted 1:1000 in blocking buffer were applied for 4 h at room temperature with agitation. Lastly, slices were washed in PBS, incubated with DAPI (Invitrogen D1306) diluted 1:50,000 in PBS, and mounted.

### Analysis of microglial morphology

Confocal z-stacks of Iba1-labelled CA1 *stratum radiatum* microglia encompassing the entire brain slice in 0.34 µm steps were imaged on a Zeiss LSM700 microscope with a Plan-Apochromat 63×/1.4 lens. Individual cells were 3D-reconstructed using the cell tracing tool Neuron Tracing v2.0 on Vaa3D (vaa3d.org), adjusting the background threshold for each image to obtain the optimal reconstruction. After checking the reconstructions against the raw images, they were analysed using custom-written MATLAB software (github.com/AttwellLab/Microglia). Briefly, based on Sholl analysis, concentric spheres were drawn at 5 µm intervals from an analyst-established cell centre (the soma is assumed to have a 5 µm radius to avoid misassigning differences in Iba1 signal within the soma as representing processes [25]). Cell process branching was profiled based on distance from the soma to assess ramification. For analysis of the effects of amyloidosis, in *App*^NL-G-F^ brain slices, both cells at 82E1-labelled Aβ plaques and >50 µm away from them were imaged. Analysis was performed with the researcher blind to genotype and Aβ signal.

### Analysis of Aβ and lysosomal burden

Immunolabelled brain slices were imaged on a Zeiss AxioScan.Z1 scanner with a Plan-Apochromat 20×/0.8 M27 lens. After background subtraction of the images by channel (using a 10-pixel rolling ball average for CD68 and Iba1 and an 80-pixel rolling ball average for Aβ) and thresholding, masks for CA1 *stratum radiatum* microglia at and >50 µm away from Aβ plaques (which were Iba1^+^/Aβ^+^ and Iba1^+^/Aβ^−^, respectively) were obtained and transposed to the binarised CD68 channel to calculate the percentage of the Iba1^+^ area that was also labelled for CD68 in microglia located at Aβ plaques and away from them.

To identify Aβ plaques in living brain tissue, a number of stains have been developed [26–28]. However, insufficient penetration into slices, labelling of only some Aβ plaque types or non-selective binding to other proteins can hinder the interpretation of results [29–31]. Thus, in this study we relied on microglial clustering around Aβ plaques (see Supplementary Fig. 1B), reported previously in *App*^NL-G-F^ mice [16, 32], as an intrinsic cue for identification of Aβ plaques (and thus of microglia at and away from them).

### Fluo-4 loading of microglia by patch-clamp electrophysiology

CA1 microglia in rat hippocampal brain slices were identified by acute labelling with isolectin-B4 coupled to Alexa-594 [25] and patch-clamped using a KCl-based intracellular solution containing (mM): 130 KCl, 4 NaCl, 10 HEPES, 0.01 BAPTA, 0.01 CaCl_2_, 10 Na-phosphocreatine, 2 MgATP, 0.5 Na_2_GTP, 0.1 Fluo-4 pentapotassium salt, adjusted to a final osmolarity of 285 ± 5 mOsm/kg and a pH of 7.2. Whole-cell recordings from microglia were obtained at a depth of > 40 μm below the slice surface using borosilicate glass pipettes with a tip resistance of 3.5–5 MΩ, resulting in access resistances of <20 MΩ that were not compensated. The average resting membrane potential of cells was −35.7 ± 2.4 mV (n=22). Throughout imaging, cells were voltage-clamped at −30 mV.

### Microglial Ca^2+^ imaging

CA1 *stratum radiatum* microglia from Cx3cr1^CreER^×GCaMP5g-IRES-tdTomato and Cx3cr1^CreER^×GCaMP5g-IRES-tdTomato × App^NL-G-F^ mice, or patch-clamped microglia filled with Fluo-4, were imaged on a Zeiss LSM780 or LSM710 two-photon microscope with a Plan-Apochromat 20×/1.0 lens and a Spectraphysics Mai Tai DeepSee eHP Ti:Sapphire infrared laser tuned to 920 nm. Fields of view of 106 µm × 106 µm were scanned with an overall acquisition time of 25 ms/frame (pixel size 0.21 µm, 1 µs pixel dwell time). Prior to longitudinal single-plane imaging (to reduce bleaching), a z-stack (encompassing the entire cell at 1 µm steps) was acquired to confirm the morphology of the imaged microglia as visualised by tdTomato.

To assess spontaneous [Ca^2+^]_i_ changes, microglial cells were imaged for 5 min (1 frame/s). Recordings were re-registered with StackReg in ImageJ/FIJI and ROIs were drawn around microglial somata or individual processes; only processes ≥1 µm in diameter were analysed to exclude filopodia [33]. In each ROI, Ca^2+^ transients were analysed with custom-written MATLAB code (github.com/AttwellLab/MyelinCalcium). A locally time-smoothened baseline (100 frames of smoothing time) was generated using a piecewise cubic Hermite interpolating polynomial fit. Ca^2+^ transients were then defined by a detection threshold for the fractional change of fluorescence (ΔF/F >2.25 × standard deviation of baseline points that had ΔF/F <0.10, to exclude contributions of Ca^2+^ transients to the baseline), confirmed using a minimal area threshold (∫ΔF/F dt >0.15), and manually checked to exclude false positives [34].

To trigger damage-evoked [Ca^2+^]_i_ changes, following a baseline recording of 20 s (1 frame/s), a focal laser lesion (6 µm radius, 177.3 µs pixel dwell time) was performed in an area >30 µm from the cell of interest, with the laser tuned to 920 nm at 80% of its maximum power. For analysis, recordings were re-registered with StackReg in ImageJ/FIJI and ΔF/F for ROIs drawn around microglial somata were calculated as (F_t_−F_o_)/F_o_, where F_t_ is the fluorescence intensity at each timepoint t and F_o_ is the average fluorescence of the pre-lesion baseline. Peak ΔF/F values were compared for statistical analysis. To help distinguish between intensity values, a multicolour (“Fire”) lookup table was applied in ImageJ/FIJI to generate the representative images provided.

### Statistics

Quantitative data are presented throughout as mean ± standard error of the mean. Normality was assessed using the D’Agostino-Pearson test. Statistical significance (defined as p <0.05) was assessed using Mann-Whitney tests (Fig. 5B–D, Suppl. Fig. 1C), two-tailed Wilcoxon matched-pairs signed rank test (Fig. 3E), one-way analysis of variance (ANOVA) followed by Dunn’s (Figs. 2D, 3B–D) or two-way ANOVA followed by Sidak’s post-hoc tests for individual comparisons (Figs. 2E, 4B, 5F). All statistical analyses were performed in Microsoft Excel 2016 and GraphPad Prism 8.

## RESULTS

### Microglia show spontaneous and damage-evoked Ca^2+^ rises *in situ*

Early studies on microglia mainly relied on cultured cells [35, 36], where gene expression is drastically changed and cells often fail to develop the complex process ramification and surveillance seen in intact microglia [14]. To overcome this limitation, while preserving the option to apply pharmacological agents in a controlled manner to modulate function, we studied microglia *in situ* in acute brain slices.

First, for an initial characterization of microglial Ca^2+^ activity in brain slices, we filled hippocampal microglia from young rats with the Ca^2+^ indicator Fluo-4 via the patch pipette to detect spontaneous Ca^2+^ transients. This experiment showed that spontaneous Ca^2+^ transients occur more frequently in microglial processes than in cell somata, with individual somata and processes showing a spontaneous frequency of events of 0.7 ± 0.4 mHz and 3.8 ± 0.7 mHz, respectively (Fig. 1A–C; see also [37] and Discussion). To study evoked Ca^2+^ responses in microglia, focal laser lesions were induced as a well-established proxy to simulate responses of microglia to brain injury, similar to focal ATP application mimicking neuronal damage [25, 38, 39]. Focal lesions lead to acute increases in microglial [Ca^2+^]_i_ at short latencies (0–5 seconds [40]). We found that in Fluo-4–filled microglia, laser lesions triggered rapid, transient Ca^2+^ signals (Fig. 1D, E).

**Fig. 1.**
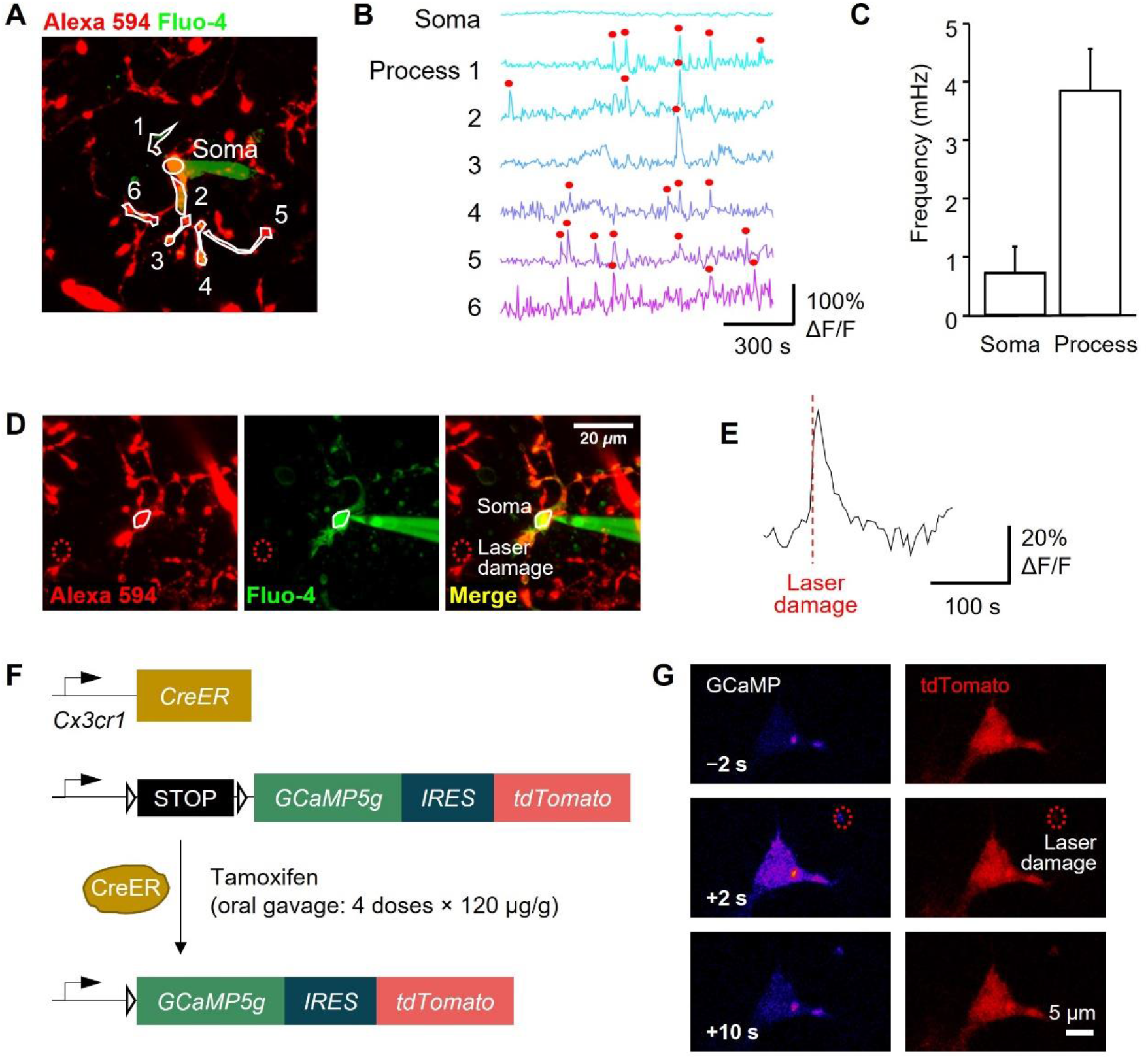
Microglia exhibit spontaneous and local damage-evoked Ca^2+^ activity in acute brain slices. **A)** Image of hippocampal slice with microglia (and vascular basement membrane) labelled with Alexa 594-conjugated isolectin B_4_, showing a patch-clamped microglial cell filled with the Ca^2+^ indicator Fluo-4. Somatic and process regions of interest (ROIs; soma and processes 1–6) are shown. **B)** Microglial [Ca^2+^]_i_ time course in the ROIs shown in (A). Red circles denote peaks detected at ≥2.5 standard deviations from the baseline. **C)** Quantification of spontaneous Ca^2+^ transient frequency, showing a higher rate in processes vs. somata (n=5 cells). **D)** As in (A) showing nearby site of laser damage (red dotted circle). **E)** A laser lesion at the site shown in (D) evokes a transient rise of [Ca^2+^]_i_. **F)** Scheme depicting the genomic changes in Cx3cr1^CreER^×GCaMP5g-IRES-tdTomato mice to drive expression of GCaMP5g and the morphological marker tdTomato in microglia induced by tamoxifen gavage. **G)** Microglia in brain slices from tamoxifen-induced Cx3cr1^CreER^×GCaMP5g-IRES-tdTomato mice show spontaneous Ca^2+^ activity (top panel), and local laser damage (red dotted circle) evokes a rapid, transient [Ca^2+^]_i_ rise in tdTomato-labelled microglia (middle and lower panels). Times indicate seconds from laser lesion.

To build on these initial findings, we employed mice that can be induced to express GCaMP5g under the control of the *Cx3cr1* promoter, which enables detection of Ca^2+^ activity in microglia (Fig. 1F) with an affinity (K_d_ 0.46 μM, ∼30-fold dynamic range [41]) similar to Fluo-4 (K_d_ 0.345 μM, ∼100-fold dynamic range [42]), and allows the Ca^2+^ concentration to be monitored in several microglia simultaneously. While use of organic Ca^2+^ indicators is very common in previous studies, these may be harmful to cells, altering their membrane potential and metabolism and causing swelling [43]. By contrast, GECIs may cause less side effects [43] and allow the study of otherwise undisturbed microglia (although all Ca^2+^ indicators will intrinsically buffer [Ca^2+^]_i_ to some extent [44]). As in young rats, we found that microglia in GCaMP5g-expressing mice showed detectable spontaneous Ca^2+^ activity that was 3.5-fold higher in rate in cell processes than in somata (as quantified below) and also exhibited local laser lesion-evoked rapid (latency ∼1 s), transient [Ca^2+^]_i_ rises (Fig. 1G).

### Proximity to Aβ plaques alters Ca^2+^ signalling in activated microglia

To test whether compartmentalised Ca^2+^ signalling is altered in brain pathology when microglia become activated, we used *App*^NL-G-F^ mice that recapitulate amyloidosis as a pathological hallmark in AD. Amyloid burden increases progressively with age in *App*^NL-G-F^ mice [16, 32]. For a reliable assessment of amyloid pathology (i.e., avoiding individual variability at onset but also allowing analysis of microglia both close to and away from plaques, which is difficult at late times because of the high Aβ plaque density), 4-month-old mice were used to study microglia in AD mice (a common age range used across studies with this model [27, 45, 46]). At this age, Aβ plaques can be detected in multiple brain regions apart from the cerebellum in perfusion-fixed slices from *App*^NL-G-F^ mice using an antibody against human Aβ (Fig. 2A, B).

**Fig. 2.**
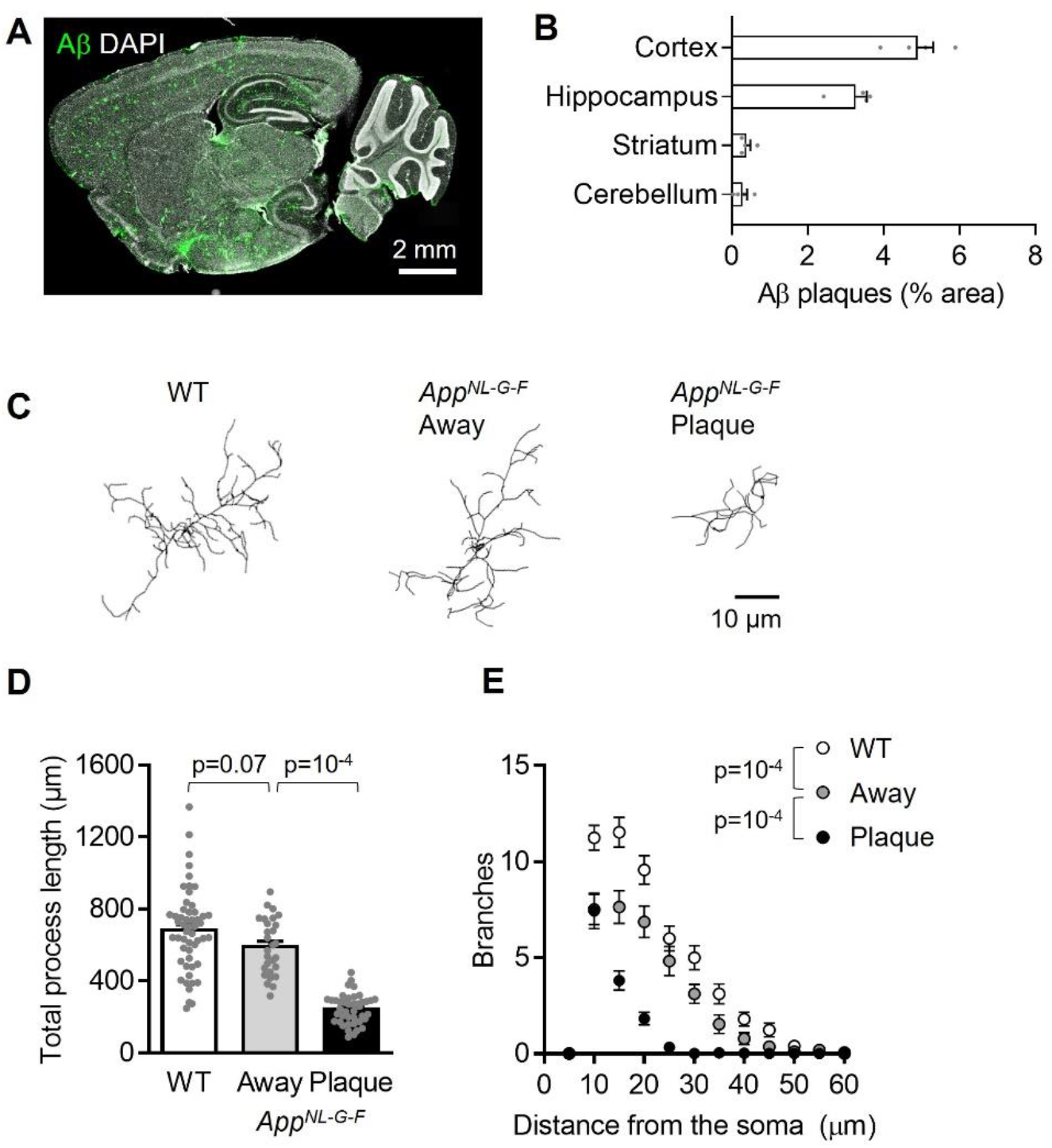
Proximity to Aβ plaques alters microglial morphology in *App*^NL-G-F^ mice. **A)** Representative sagittal brain slice from a 4-month-old *App*^NL-G-F^ mouse showing Aβ plaques (82E1 antibody, green) and cell nuclei (DAPI, white) for reference. **B)** Quantification of the area covered by Aβ plaques in the neocortex (layer 2/3), dorsal hippocampus (CA1), cerebellum (molecular layer) and dorsal striatum of 4 *App*^NL-G-F^ mice. **C)** Representative 3D-reconstructed *App* wild-type (WT) microglia, and *App*^NL-G-F^ microglia at or >50 µm away from Aβ plaques in 4-month-old mice. **D–E)** Sholl analysis-derived (D) total process length and (E) number of process branches at 5 μm increment distances from the soma, showing a sharp deramification of microglia at plaques (n=53 *App* WT cells, 28 *App*^NL-G-F^ cells away from plaques and 42 *App*^NL-G-F^ cells at plaques from 3 mice each).

We first assessed whether microglial morphology was affected by proximity to Aβ plaques, as has been shown previously in other non-knock-in models of AD, which have a differing development of disease parameters [47–49]. While microglia were highly ramified in wild-type mice, their overall process length and ramification were reduced in *App*^NL-G-F^ mice (Fig. 2C–E). The total process length per cell was reduced by 13.4% in *App*^NL-G-F^ microglia >50 μm away from Aβ plaques compared with wild-type microglia (not significantly different, p=0.07) and by 64.3% in *App*^NL-G-F^ microglia at Aβ plaques compared with wild-type microglia (p=10^-4^; Fig. 2D). Similarly, process ramification was reduced closer to Aβ plaques, with *App*^NL-G-F^ microglia showing less branched processes and less branches near the soma compared with wild-type cells (Fig. 2E).

Microglia are known to readily phagocytose Aβ debris [50]. In our *App*^NL-G-F^ model, engulfment of Aβ inside microglial CD68-positive lysosomes was confirmed by confocal imaging (Supplementary Fig. 1A). CD68 coverage approximately doubled in hippocampal microglia at Aβ plaques compared with cells away from plaques (p=10^-4^; Supplementary Fig. 1B, C), suggesting an increase in the phagocytic capacity of microglia in the presence of Aβ. This is consistent with the increase in CD68 expression reported in plaque-proximal microglia from other AD models and in microglia from patients with AD [51–53].

Together, these results suggest that microglial responses to Aβ depend considerably on the cells’ local environment, whereby the cells closer to Aβ plaques exhibit stronger changes in morphology and lysosomal content while those further away display an intermediate phenotype closer to wild-type microglia. Therefore, in subsequent experiments we analysed data from cells at Aβ plaques and away from plaques separately (using a distance threshold of >50 µm from the nearest plaque edge as the defining criterion for plaque-distant microglia).

We next examined whether the microglial morphological changes in the *App*^NL-G-F^ mice are reflected in a specific Ca^2+^ activity pattern. We found that the frequency of spontaneous Ca^2+^ transients in microglia somata was dependent on Aβ pathology and proximity to plaques (Fig. 3A, B). While somata of wild-type microglia exhibited Ca^2+^ transients at a frequency of 1.5 ± 0.5 mHz, *App*^NL-G-F^ microglia located away from Aβ plaques and at Aβ plaques had somatic transient rates of 4.2 ± 0.8 mHz (not significantly different from wild-type, p=0.1) and 7.3 ± 1.0 mHz (p=10^-4^), respectively. Of note, this effect was not seen in microglial processes, where spontaneous Ca^2+^ events occurred independently of *App*^NL-G-F^ genotype; the transient rate was 9.1 ± 2.1 mHz per process in wild-type microglia, 9.7 ± 1.2 mHz in *App*^NL-G-F^ microglia away from Aβ plaques (p=0.9) and 8.8 ± 0.8 mHz in plaque-associated cells (p=0.9; Fig. 3C, D).

**Fig. 3.**
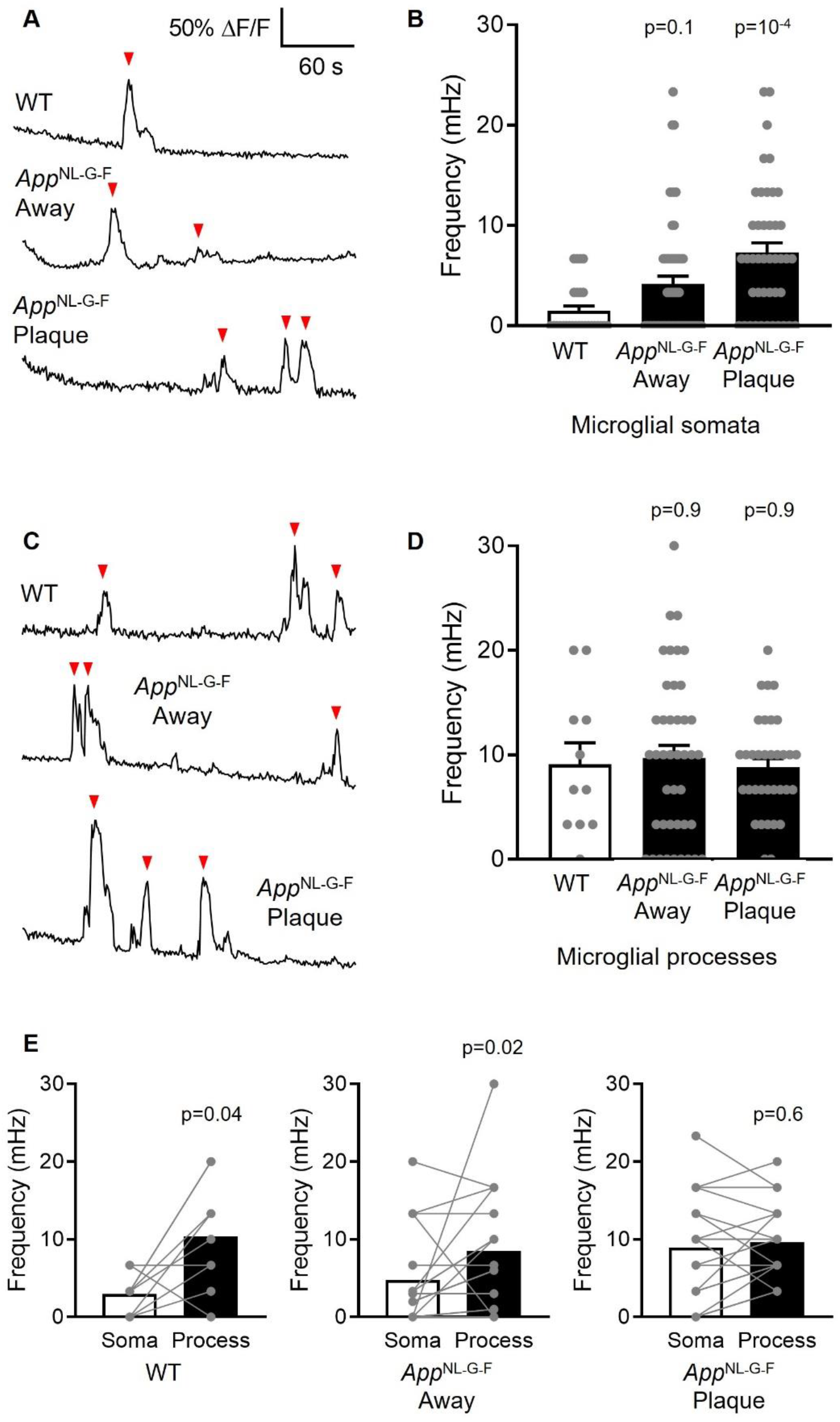
Spontaneous Ca^2+^ activity in microglial somata is increased near Aβ plaques. **A)** Representative traces showing spontaneous Ca^2+^ transients (red arrowheads) in microglial somata from *App* wild-type (WT) and *App*^NL-G-F^ mice at, or >50 µm away from, Aβ plaques. **B)** Quantification of spontaneous Ca^2+^ transient frequency, showing a higher rate near Aβ plaques (*App* WT: n=27 somata from 5 mice; *App*^NL-G-F^ away from plaques: n=56 somata from 8 mice; *App*^NL-G-F^ at plaques: n=43 somata from 8 mice). **C)** As (A), but for processes. **D)** As (B), but for processes, showing no effect of Aβ status (*App* WT: n=11 processes from 3 mice; *App*^NL-G-F^ away from plaques: n=42 processes from 6 mice; *App*^NL-G-F^ at plaques: n=34 processes from 6 mice). **E)** Paired comparisons between process and somatic Ca^2+^ transient rate, showing that, for a given microglial cell, spontaneous Ca^2+^ activity per process was higher in processes in *App* WT (n=9 cells from 3 mice) and *App*^NL-G-F^ microglia away from plaques (n=18 cells from 6 mice), but not in microglia at Aβ plaques (n=19 cells from 6 mice) where the somatic frequency is similar to the frequency per process in WT mice.

Because of the different Ca^2+^ activity in microglial somata vs. processes when averaged across cells, we also analysed a random subset of cells calculating their individual transient rates in the somata and processes in a paired manner (averaging over processes to obtain a single mean process value for each cell: Fig. 3E). In nearly all cells, microglial processes displayed a significantly higher rate of Ca^2+^ transients per process than somata in wild-type mice (3.5-fold higher; p=0.04), which was also seen in microglia away from Aβ plaques in *App*^NL-G-F^ mice (1.8-fold higher; p=0.02). In contrast, microglia at Aβ plaques showed a similar rate of spontaneous Ca^2+^ events in their processes and somata (1.1-fold higher, p=0.6).

Next, we studied whether amyloid pathology altered lesion-evoked Ca^2+^ responses in microglia. While laser injury triggered a robust, rapid transient rise of [Ca^2+^]_i_ in wild-type microglial somata, this response was impaired in the *App*^NL-G-F^ mice (Fig. 4). In *App*^NL-G-F^ microglia away from Aβ plaques, the peak Ca^2+^ response was reduced by 31.2% (p=0.09), and in those at Aβ plaques the peak Ca^2+^ response was reduced by 77.9% (p=10^-4^). Together, these data suggest that, while Aβ increases spontaneous Ca^2+^ activity in microglial somata, it renders microglia less able to respond to substances (such as ATP and ATP-derived metabolites) released acutely by laser-evoked brain damage. In both cases, microglia >50 μm distant from Aβ plaques exhibited an intermediate phenotype between those at plaques and their wild-type counterparts.

**Fig. 4.**
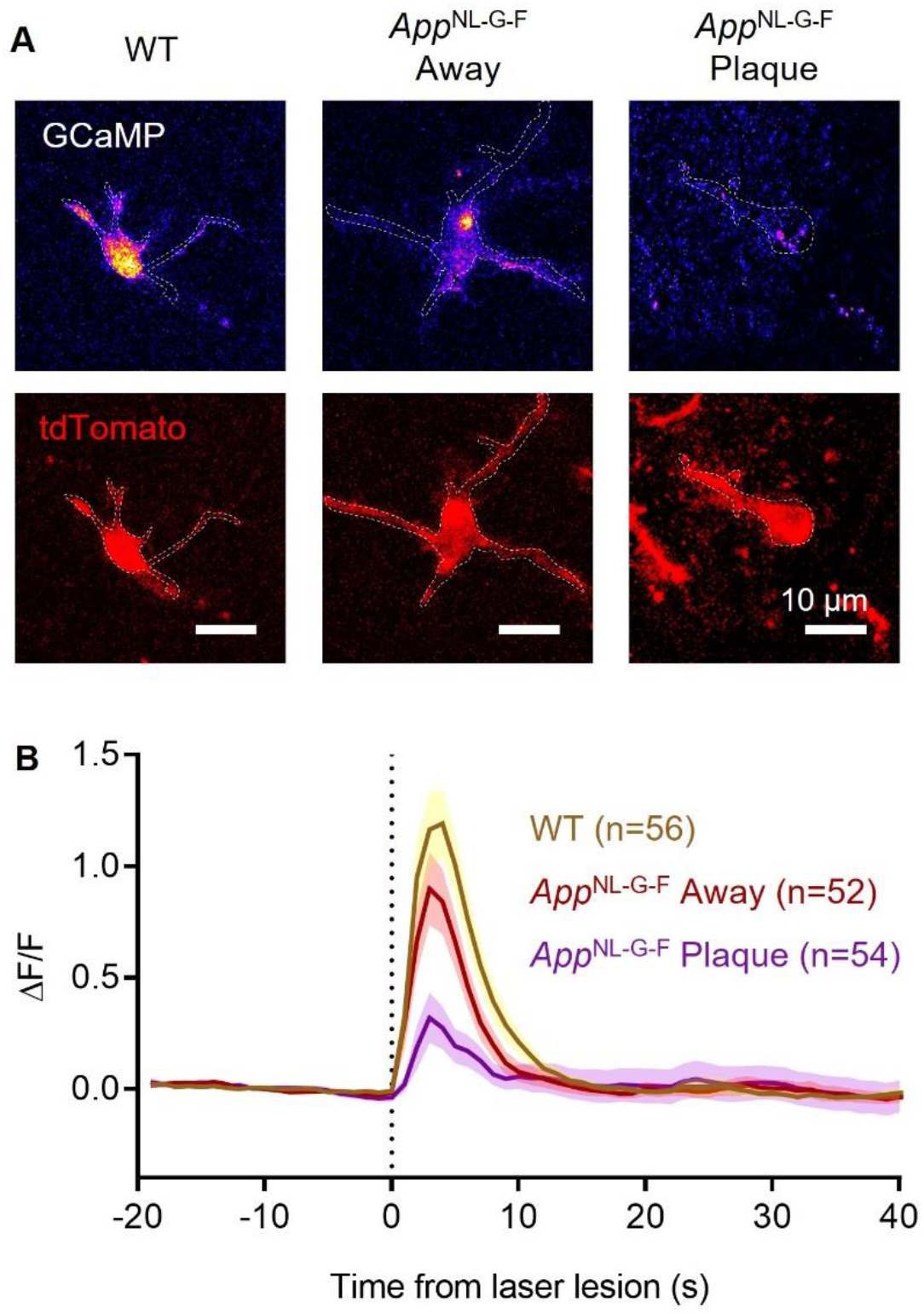
Damage-evoked Ca^2+^ rises are reduced in microglia near Aβ plaques. **A)** Representative microglia (tdTomato, red, lower panels) from *App* wild-type (WT) and *App*^NL-G-F^ mice at Aβ plaques or away (>50 µm) from plaques. Laser damage-evoked Ca^2+^ rises (upper panels, shown on a “Fire” scale) are measured by GCaMP5g fluorescence changes (ΔF/F shown at peak). **B)** Quantification of somatic [Ca^2+^]_i_ levels over time (ΔF/F) showing their increase upon laser lesion (vertical dashed line). *App*^NL-G-F^ microglia, both at (n=54 from 8 mice; p= 10^-4^) and away from Aβ plaques (n=52 from 8 mice; p=0.03), showed a significantly reduced damage-evoked [Ca^2+^]_i_ rise compared with *App* WT microglia (n=56 from 5 mice).

### Ageing alters microglial Ca^2+^ signalling differently from Aβ pathology

There is growing interest in the cellular processes of ageing and how these relate to neurodegenerative processes [22, 54–56]. Therefore, we analysed whether the changes in Ca^2+^ signalling that we found in microglia from AD mice may also occur in old (P300–310) wild-type mice.

While the frequency of spontaneous Ca^2+^ transients in old microglial somata was not significantly changed with age (40.0% lower than in microglia from young adult (P120-130) mice but not statistically significantly different; p=0.46), microglial processes from old mice exhibited a significantly reduced frequency of Ca^2+^ transients (70.7% lower; p=10^-3^; Fig. 5A– D). We also analysed lesion-evoked somatic Ca^2+^ responses and found that they were largely abolished in microglia from old mice, with the peak Ca^2+^ response reduced by 77.4% (p=10^-4^) compared with microglia from young adult animals (Fig. 5E, F). These data suggest that normal ageing affects microglial Ca^2+^ activity differently from the pathological conditions of amyloidosis, reducing not only brain damage-evoked but also spontaneous Ca^2+^ activity in the cell processes.

**Fig. 5.**
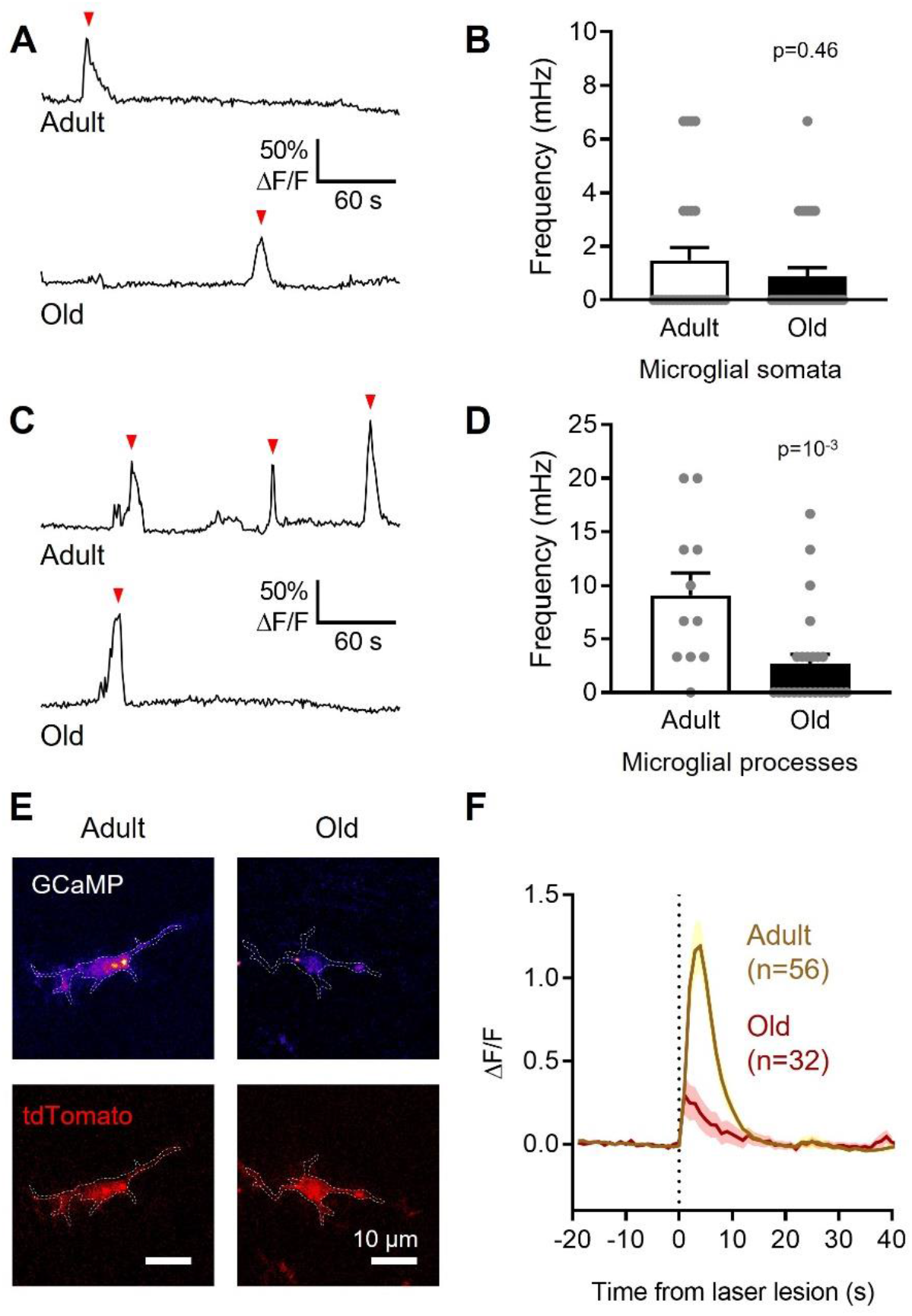
Microglia from old mice exhibit reduced spontaneous and damage-evoked Ca^2+^ activity compared with microglia from young adult mice. **A)** Representative traces showing spontaneous Ca^2+^ transients (red arrowheads) in microglial somata from P120–130 (young adult) and P300–310 (old) mice. **B)** Quantification of spontaneous Ca^2+^ transient frequency, showing no difference between somata of microglia from young adult and old mice (n=27 somata from 5 young adult mice and 30 somata from 3 old mice). **C)** As (A), but for processes. **D)** As (B), but for processes, showing a statistically significant decrease in spontaneous Ca^2+^ transient frequency with age (n=11 processes from 3 young adult mice and 25 processes from 3 old mice). Data from *App* wild-type microglia in Fig. 3 are included as young adult microglia for comparison. **E)** Representative microglia (tdTomato, red) from P120–130 (young adult) and P300–310 (old) mice. Laser damage-evoked Ca^2+^ rises are measured by GCaMP5g fluorescence changes (ΔF/F shown at peak, “Fire” scale). **F)** Quantification of somatic [Ca^2+^]_i_ levels over time (ΔF/F) showing their increase upon laser lesion (vertical dashed line). Microglia from old mice (n=32 cells from 3 mice) showed a significantly reduced damage-evoked [Ca^2+^]_i_ rise compared with microglia from young adult mice (n=56 from 5 mice; p=10^-4^). Data from *App* wild-type microglia in Fig. 4 are included as young adult microglia for comparison.

## DISCUSSION

Intracellular Ca^2+^ elevations are commonly considered an indicator of cellular activity in neurons and astrocytes; however, knowledge of microglial Ca^2+^ signalling is still sparse despite its importance in controlling key cellular functions. While spontaneous Ca^2+^ transients are extremely infrequent in acutely isolated or cultured microglia from adult mice [13], *in vivo* experiments on OGB-1–electroporated microglia showed that ∼20% of them displayed transients over 15 minutes [40], and a later study using GECI-expressing microglia found that ∼4% of cells were active over 20 min [7]. This suggests that the environment, type of preparation and Ca^2+^ sensor, and the activation state may all affect spontaneous Ca^2+^ activity in microglia. We found that the great majority of Ca^2+^ transients in healthy rat or mice microglia occurred in the cells’ processes, whether studied with the soluble Ca^2+^ sensor Fluo-4 or with GCaMP5g. This finding, which is in agreement with previous work using GECI-expressing mice [37], implies that microglial Ca^2+^ signalling is finely regulated across cell compartments (as in astrocytes [57]). Ca^2+^ transients may fulfil different functions in the soma vs. processes, which may be related to differential expression and localisation of Ca^2+^ signal-transducing membrane receptors and intracellular Ca^2+^-dependent organelles, such as the endoplasmic reticulum or the endolysosomal compartment.

Although microglia rarely generate Ca^2+^ transients in the healthy brain, elevations in [Ca^2+^]_i_ are much more prominent under pathological conditions such as epileptiform activity, acute brain injury, neuroinflammation, and neurodegeneration [7, 9, 15, 37, 40]. In AD, the comparatively well characterised changes in microglial morphology and biomarker expression contrast with the still poorly understood changes of Ca^2+^ signalling in these cells, and how these vary between plaque and plaque-remote regions and in different parts of each microglial cell. In *App*^NL-G-F^ knock-in mice, we observed that microglial deramification and increased lysosomal burden depend critically on the cells’ proximity to Aβ plaques. This is consistent with studies of transgenic models of AD, showing that changes in microglial gene expression, morphology and electrophysiological properties [32, 48, 58], as well as synapse loss [59], occur predominantly near plaques. In agreement with this, our study revealed significant differences in spontaneous and damage-evoked Ca^2+^ responses in the *App*^NL-G-F^ model of AD between Aβ plaque-proximal and plaque-distant microglia, with the latter presenting an intermediate phenotype closer to that of wild-type cells. Notably, and adding to previous work, we found that these changes were also dependent on the subcellular compartment, revealing differential effects of Aβ on Ca^2+^ activity in processes compared with somata. While the Ca^2+^ activity of microglia has been previously studied in mice modelling AD [60], this was investigated with two different transgenic mouse strains pooled together (both overexpressing *App*) and using a dye indicator, OGB-1, electroporated into cortical microglial cells to estimate changes in [Ca^2+^]_i_. Here, we took advantage of GCaMP5g to visualise changes in [Ca^2+^]_i_, because intrinsic GECI expression may affect cell function less than using patch-clamping or electroporation to introduce a dye into the cell (although the affinity and dynamic range of the indicator used will also affect its usefulness).

We found that proximity to Aβ plaques increased the rate of spontaneous Ca^2+^ transients in the somata of hippocampal microglia. This is in line with results from Brawek et al. (2014), noting an increase in the fraction of microglia with Ca^2+^ activity in older, transgenic amyloid-depositing mice (although that study, which analysed cortical rather than hippocampal microglia, did not observe any changes in the rate of spontaneous Ca^2+^ transients in the cells that exhibited them [60]). Increased Ca^2+^ activity in microglial cells exposed to Aβ may be due to Aβ-evoked UTP/UDP release by stressed neurons acting on microglial metabotropic P2Y_6_ receptors [61] or Aβ-mediated mechanotransduction mediated via Piezo1 Ca^2+^-permeable ion channels involved in amyloid clearance [10, 62]. However, while the Ca^2+^ transient rate in microglial somata depended on their proximity to Aβ plaques, the higher rate in microglial processes did not. Given the decrease in ramification of microglia at Aβ plaques, the increased Ca^2+^ frequency in somata vs. processes could also reflect the membrane and Ca^2+^ stores of the latter being pulled back into the soma, or a redistribution of receptors underlying Ca^2+^ responses [60]. Of note, the higher Ca^2+^ activity in the soma of microglia at Aβ plaques correlates with a large increase in CD68 immunoreactivity in these cells, consistent with an increased phagocytic rate (which has been linked to microglial Ca2+ activity [9]). Indeed, in microglia this marker of mature phagocytic lysosomes is localised mainly in somatic regions [63], where it may support an increased Ca^2+^-dependent phagocytic activity in plaque-associated microglia.

Another key finding from the present study is that microglia at Aβ plaques generated reduced responses to laser lesions. This suggests that cells activated by Aβ might be less able to mount large Ca^2+^ responses even though they exhibited an increased baseline of spontaneous transients in the absence of acute injury. In support of this hypothesis, basal Ca^2+^ levels are higher in microglia isolated from post-mortem AD brains than in cells from non-dementing donors, but their ATP-evoked responses are smaller [64]. Similarly, in *App*-overexpressing transgenic mice, chemotaxis towards local ATP is also impaired in microglia near Aβ plaques [60], suggesting that injury-evoked Ca^2+^ rises might also be reduced in these cells. The decreased response to laser damage–evoked nucleotide release may reflect reduced expression of P2Y_12_ receptors as a result of microglial activation [65, 66].

Finally, it is important to assess how changes in Ca^2+^ responses in AD compare to other conditions, as diverse stimuli inducing microglial activation might converge by regulating [Ca^2+^]_i_ as a core signalling element. The microglial genetic signature is similar between AD and old age [22, 54–56], and accumulation of Aβ (and tau) in the disease are thought to exacerbate processes involved in ageing, such as cellular senescence, Ca^2+^ dyshomeostasis, and inflammation [67]. Of note, our study revealed that amyloidosis and ageing affected the Ca^2+^ activity of hippocampal microglia differently. In AD mice, spontaneous Ca^2+^ activity was increased in the somata of microglia but not in their processes, compared with age-matched, non-AD mice. By contrast, spontaneous Ca^2+^ activity in aged mice was unchanged in microglial somata, but decreased in processes, compared with younger adults. Interestingly and in line with the lack of observed changes in somatic Ca^2+^ activity with age, microglial CD68 expression is only mildly altered between young and old mice [68, 69]. Of note, Brawek et al. (2014) suggested that the fraction of microglia showing spontaneous Ca^2+^ activity may be increased in ageing [60], and a subsequent study revealed a bell-shaped pattern with mice of 9–11 months of age (i.e., what we refer to as old age) showing a higher rate of Ca^2+^ transients than both 2–4- and 18–21-month-old animals [70]. This apparent difference with our results may arise from differences between the AD mouse models used, between the brain regions studied (neocortex vs. hippocampus) or in the Ca^2+^ detection method used (dye electroporation vs. GECI). Although each method has its own limitations, the high sensitivity of microglia to mechanical influences [62, 71], with differences reported even between *in vivo* preparations (acute vs. chronic window preparations [37]), may be a critical factor to consider along with an age-dependent increase in cellular vulnerability and thus a reduced experimental resilience to mechanical perturbations. While reduced lesion-evoked Ca^2+^ responses are common to microglia in both physiological ageing and AD, our study suggests that clear differences exist at the subcellular level for spontaneous Ca^2+^ events, the details of which should be clarified by future work. Overall, our work highlights the multifaceted functional states that microglia can adopt depending on their environment (i.e., in a healthy brain, in AD whether located at Aβ plaques or away from them, and in the aged brain), and shows that Ca^2+^ activity is differently regulated on a subcellular level in each case.

## DATA AVAILABILITY

All code used for analysing microglial morphological parameters and Ca^2+^ imaging data has been deposited in GitHub (github.com/AttwellLab). All other study data are included in the article and/or Supplementary Information.

## Acknowledgements

This work was supported by European Research Council (BrainEnergy) and Wellcome Investigator Awards (099222) to DA, a Wellcome Trust four-year PhD studentship to PI and an Alzheimer Forschung Initiative grant (21072) to CM. For the purpose of open access, the authors have applied a CC-BY public licence to any Author Accepted Manuscript version arising from this submission.

---

**Supplementary Fig. 1.**
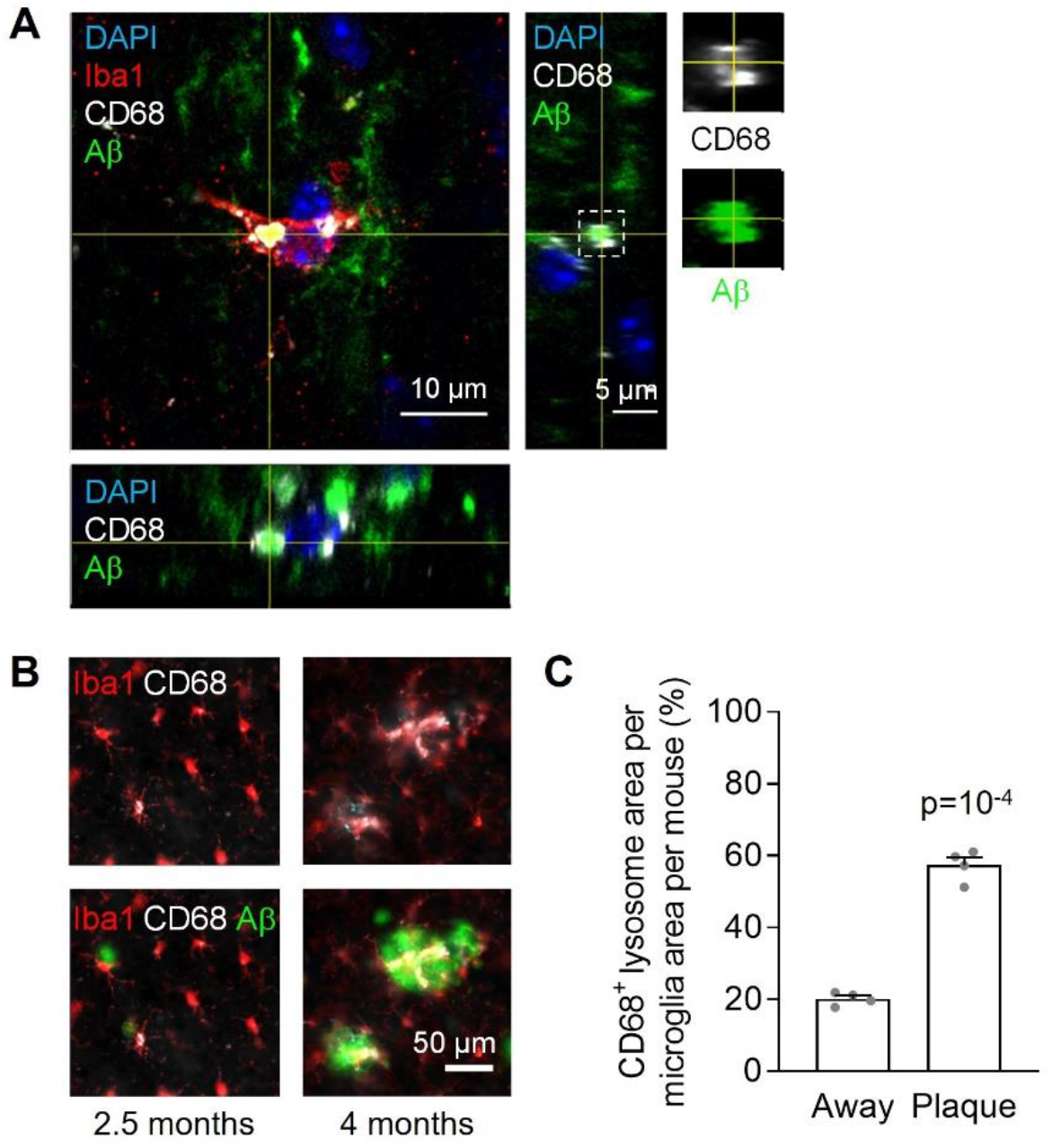
Proximity to Aβ plaques increases the lysosomal burden in microglia from *App*^NL-G-F^ mice. **A)** Microglial cell with phagocytosed Aβ in a 4-month-old *App*^NL-G-F^ mouse. Orthogonal projections at the level of the crosshairs show internalisation of Aβ by the microglial cell (intensity of Aβ channel is adjusted to see internalised Aβ more clearly). Right panels show the indicated area (white dashed square) at higher magnification to highlight the presence of Aβ within the CD68^+^ lysosome. **B)** Representative images of hippocampal microglia (Iba1, red) in *App*^NL-G-F^ mice at 2.5 months (left) and 4 months of age (right). At 4 months, when large Aβ plaques (green) have already developed, microglia show increased expression of the lysosomal marker CD68 (white) as they cluster around them. **C)** Quantification of the fraction of the microglial area covered by CD68 in the hippocampus of four 4-month-old *App*^NL-G-F^ mice, showing a sharp increase in plaque microglia compared to cells away from plaques.

